# A machine learning model for disease risk prediction by integrating genetic and non-genetic factors

**DOI:** 10.1101/2022.08.22.504882

**Authors:** Yu Xu, Chonghao Wang, Zeming Li, Yunpeng Cai, Ouzhou Young, Aiping Lyu, Lu Zhang

## Abstract

Polygenic risk score (PRS) has been widely used to identify the high-risk individuals from the general population, which would be helpful for disease prevention and early treatment. Many methods have been developed to calculate PRS by weighted aggregating the phenotype-associated risk alleles from genome-wide association studies. However, only considering genetic effects may not be sufficient for risk prediction because the disease risk is not only related to genetic factors but also non-genetic factors, e.g., diet, physical exercise et al. But it is still a challenge to integrate these genetic and non-genetic factors into a unified machine learning framework for disease risk prediction. In this paper, we proposed PRSIMD (PRS Integrating Multi-source Data), a machine learning model that applies posterior regularization to integrate genetic and non-genetic factors to improve disease risk prediction. Also, we applied Mendelian Randomization analysis to identify the causal non-genetic risk factors for the selected diseases. We applied PRSIMD to predict type 2 diabetes and coronary artery disease from UK Biobank and observed that PRSIMD was significantly better than the methods to calculate PRS including *p*-value threshold (P+T), PRSice2, SBLUP, DMSLMM, and LDpred2. In addition, we observed that PRSIMD achieved the better predictive power than the composite risk score.

## I. Introduction

Polygenic risk score (PRS) is an approach to predict an individual’s risk of particular phenotypes using genetic variants, such as single nucleotide polymorphisms (SNPs). It is usually calculated by weighted (e.g., effect size) aggregating an individual’s risk alleles identified by genome-wide association studies (GWAS) [1]. Many computational models have been proposed to calculate PRS, which can be classified into three categories [2]: 1. thresholding and pruning methods, such as P+T [3] and PRSice2 [4]; 2. penalized regression methods, such as SBLUP [5] and DBSLMM [6]; and 3. Bayesian methods such as LDpred2 [7] and PRS-CS [8]. Recent large-scale GWAS [9], [10] provide us an unprecedented opportunity to identify the susceptible disease variants and apply PRS to identify the high-risk individuals from general populations [11], [12].

However, the risk of most diseases would also be influenced by non-genetic factors. Several previous studies considered both genetic and non-genetic factors for disease risk prediction. O’Sullivan et al. [13] implemented *”CHA*_*2*_*DS*_*2*_*-V ASc”* model to integrate PRS and eight clinical factors (Congestive heart failure, Age>75, Diabetes Mellitus, etc.) in a logistic regression to predict the risk of atrial fibrillation stroke. Yiwey et al. [14] selected 83 SNPs to calculate PRS and incorporated it with the 5-year risk probability from the breast cancer surveillance consortium to build the risk model. Moldovan et al. [15] applied the composite risk score (CRS) to predict the risk of type 2 diabetes. CRS is defined as the weighted sum of three risk measures: 1. PRS; 2. age; and 3. individual characteristics (body mass index (BMI), birth weight, or comparative body size at age ten). The weights of these three measures were learned from the training set. However, these studies were mainly designed for particular diseases, and the integration of genetic and different types of non-genetic factors still challenges disease risk prediction.

We proposed a machine learning model called PRSIMD (Polygenic Risk Score Integrating Multi-source Data) to integrate the genetic and non-genetic factors using posterior regularization [16], [17] for disease risk prediction. PRSIMD converts the PRS into posterior distribution using logistic regression and adopts log-linear model to aggregate non-genetic factors into prior knowledge distribution. PRSIMD employs Kullback-Leibler (KL) divergence as an additional loss to force the posterior distribution to approach the prior knowledge distribution in the training procedure.

We compared PRSIMD with five algorithms to predict PRS including *p*-value threshold (P+T), PRSice2, SBLUP, DMSLMM, and LDpred2; and the composite risk score (CRS) by integrating PRS and non-genetic factors on coronary artery disease (CAD) and type 2 diabetes (T2D) (**Fig. 1A**) from UK Biobank (UKBB) [18]. For each disease, we selected the top 30 most significant disease causal risk factors (**Fig. 1C and 2**) in physical measures and lifestyles using Mendelian Randomization (MR) [19], which combined with age and sex as the risk factors for the diseases. The experimental results showed that PRSIMD outperformed all PRS algorithms that only involved genetic factors, with about 24% average increase of AUROC (area under the receiver operating characteristics) on CAD and T2D compared with the second-best method (**Fig. 3A-B**). We also observed that PRSIMD had a better performance than CRS (**Fig. 3C-D**). PRSIMD increased the AUROC at least 0.013 (0.035) on CAD (T2D) compared to CRS.

**Fig. 1.**
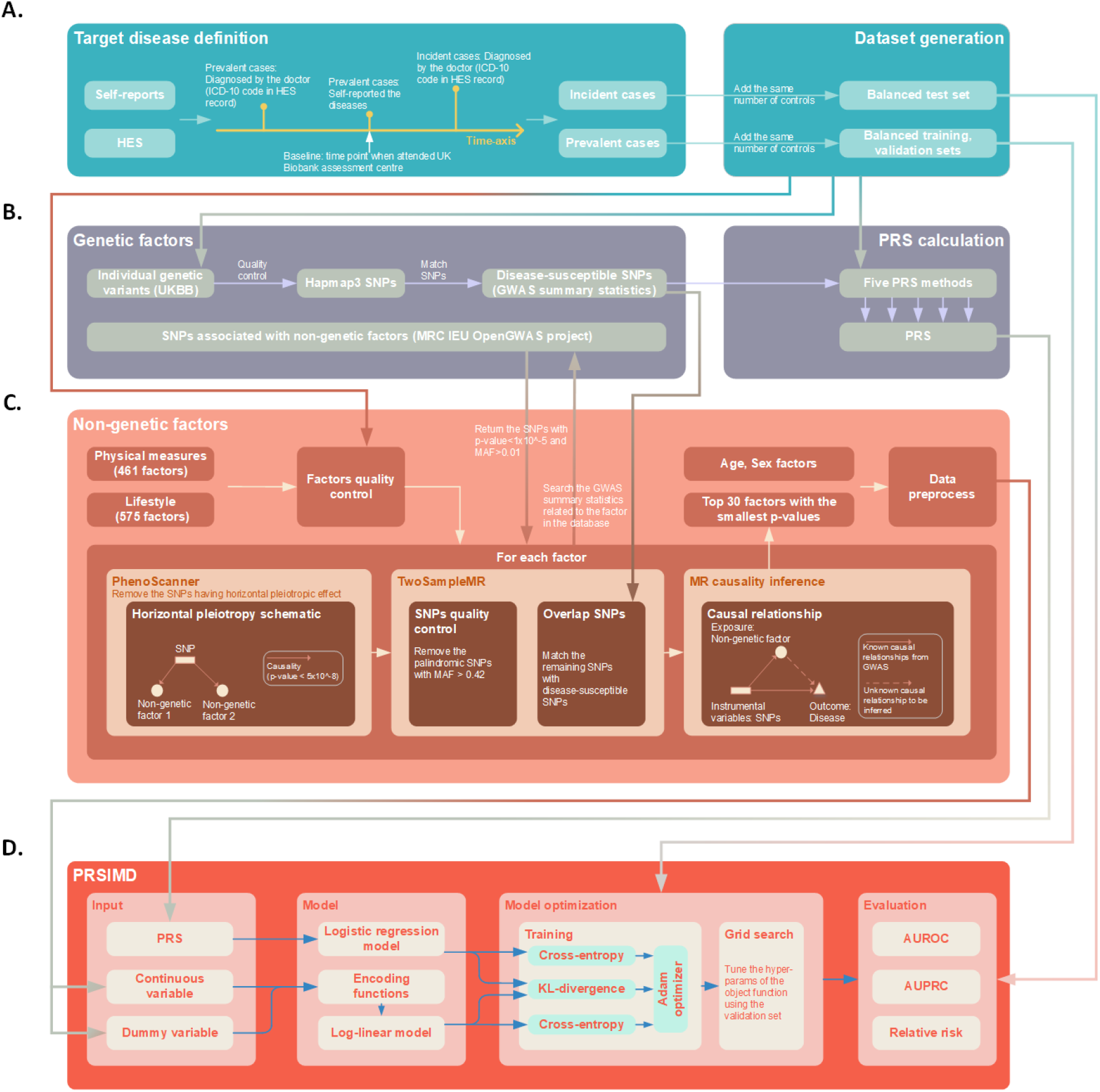
An overview of the workflow. **A**. We defined CAD and T2D in UK biobank using self-reports and hospital episode information. The prevalent (incident) cases were used to generate the training and validation (test) sets. **B**. We used the disease-susceptible SNPs to calculate PRS by five existing PRS methods. The SNPs associated with the non-genetic factors from OpenGWAS project were used for Mendelian Randomization analysis. **C**. We utilized PhenoScanner and TwoSampleMR to construct the instrumental variable associated with the factor and the disease. The instrumental variable was used to infer the potential causality between the factor and the disease. **D**. The input of PRSIMD consists of the PRS and the selected non-genetic risk factors. We applied posterior regularization to train the model and then evaluated it by using three metrics.

## II. Method

### A. Target disease definitions and datasets generation

We defined CAD and T2D based on self-reports and ICD-10 codes in hospital episode statistics (HES). We generated a mapping table to match the ICD-10 codes with participants’ self-reported diseases based on the descriptions of their symptoms: 1. CAD: I21.X, I22.X, I23.X, I241, and I252; 2. T2D: E11.X. X is a wildcard [20]. As shown in the left panel of **Fig. 1A**, the prevalent cases were defined as participants having self-reported diseases or corresponding ICD-10 codes that were diagnosed before the baseline. The incident cases were defined as disease events occurring after the baseline (among participants who did not have the same prior diseases). The participants used in this paper were restricted to the white British without kinship to each other.

We generated a general control pool for participants who were never diagnosed with any of the two target diseases. The controls were selected randomly and combined together with prevalent and incident cases. We used two-thirds of the prevalent cases and controls to generate the training sets, and the remaining one-third to generate the validation sets. The controls in each disease are non-overlapping. We collected incident cases and controls as test sets to investigate the predictive power of PRSIMD. We used the same number of cases and controls for each disease to ensure that the datasets were balanced (the right panel of **Fig. 1A**).

### B. Data usage of genetic and non-genetic factors

For genetic factors, we selected the high-quality SNPs from UKBB by matching them with HapMap3 (used in PRS-CS [8]). Their *p*-values and effect sizes were further matched to the GWAS summary statistics of CAD and T2D from previous publications (**Fig. 1B** and **Table. I**).

**TABLE I.**
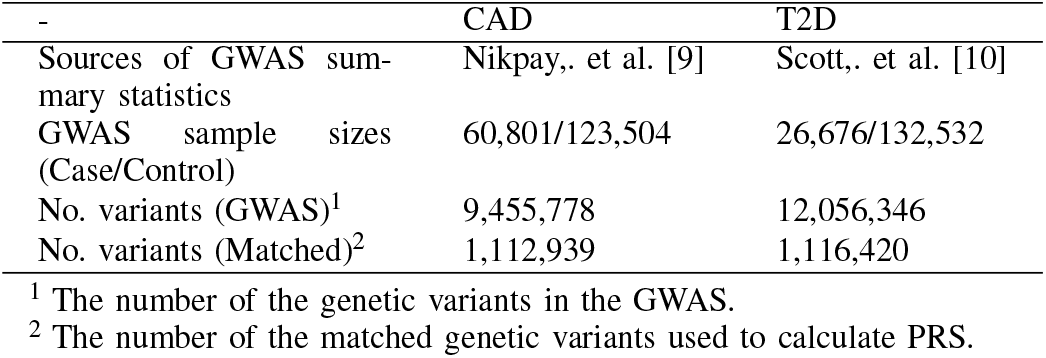
GWAS summary statistics for CAD and T2D

|  | CAD | T2D |
| --- | --- | --- |
| Sources of GWAS summary statistics | Nikpay, et al. [9] | Scott, et al. [10] |
| GWAS sample sizes (Case/Control) | 60,801/123,504 | 26,676/132,532 |
| No. variants (GWAS)<sup>1</sup> | 9,455,778 | 12,056,346 |
| No. variants (Matched)<sup>2</sup> | 1,112,939 | 1,116,420 |

<sup>1</sup> The number of the genetic variants in the GWAS.
<sup>2</sup> The number of the matched genetic variants used to calculate PRS.

For non-genetic factors, we selected the disease risk non-genetic factors from physical measures and lifestyles in UKBB by fitting logistic regressions in the training sets and filtered out the non-informative factors if 1. *p*-value ≥ 0.05; and 2. over 50,000 missing values (the upper-left panel of **Fig. 1C**). We then constructed the instrumental variable for each selected non-genetic factor (denoted as *k*_*i*_; *i* = { 1, 2, …, *n* } ; n represents the number of the candidate non-genetic factors) to infer its causality to the disease. Firstly, we used the GWAS summary statistics of k_*i*_ from the MRC IEU OpenGWAS project [21], [22] to select the SNPs if 1. they had *p*-value< 1 × 10^*−*5^; 2. minor allele frequencies (MAFs) > 0.01 [23]. To avoid horizontal pleiotropy, the SNPs were removed if they were significantly associated with the other confounding non-genetic factors (*p*-value< 5 × 10^*−*8^) by utilizing PhenoScanner [24] (the lower-left panel of **Fig. 1C**). Because the palindromic SNPs (having A/T or G/C alleles) with MAF > 0.42 were regarded as not inferrable [23], we applied TwoSampleMR [25] to remove them and match the remaining SNPs with the disease-susceptible SNPs [10], [26], resulting in the SNPs associated with both *k*_*i*_ and the diasease (the lower-middle panel of **Fig. 1C**). The matched SNPs were used as the instrumental variable of *k*_*i*_. Lastly, we performed Inverse Variance-Weighted method [19] to infer the causalities of non-genetic factors (*k*_*i*_) and the two diseases, respectively (the lower-right panel of **Fig. 1C**). The top 30 causal non-genetic factors with the smallest *p*-values were selected and combined with age (when attended UKBB assessment centre) and sex as the risk factors for the disease. Moreover, we applied the median value (for continuous factors) and the most frequent value (for integer and categorical factors) to impute the missing values in the data (the upper right panel of **Fig. 1C**).

Besides, we applied L1 penalized logistic regressions to estimate the effect sizes and 95% confidence intervals (CIs) of the selected non-genetic factors on the risk of CAD and T2D. For categorical factors, we first selected an item from the factor as a baseline, and the ORs (odd ratio) of the other items in this factor were calculated by comparing them with the baseline. In addition, we also included age and sex as covariates in the logistic regression.

### C. Modelling the genetic factors

We calculated the PRSs using P+T [3], PRSice2 [4], SBLUP [5], DBSLMM [6], and LDpred2 [7] using the summary statistics from previous publications [9], [10]. The PRS for each disease was transformed into a probabilistic distribution (*P(y*|*x*_*s*_; *θ*)) by logistic regression:

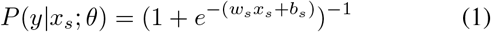

where *x*_*s*_ represents the value of the PRS, and *y* is the disease label (1 for the affected patients, 0 for the healthy controls). *θ* = { *w*_*s*_, *b*_*s*_ *}* represent the weights that can be learned from the training set. We used cross-entropy [27] as loss function for model optimization.

### D. Modelling the non-genetic factors

We considered the integer (e.g., age) and continuous (e.g., BMI) non-genetic factors as continuous variables. In contrast, we transformed the binary (e.g., sex) and multiple categorical (e.g., cheese intake) non-genetic factors using one-hot encoding. Furthermore, we designed a set of encoding functions (denoted as *ϕ)* for the non-genetic factors to weight and rescale their values into [-1,1] using the non-linear activation function *tanh*.

We assigned a learnable scalar w to weight the continuous variable. For instance, age and BMI are the most common continuous variables, which can be encoded by the function of *ϕ*_*age*_ and *ϕ*_*bmi*_:

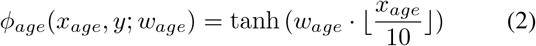

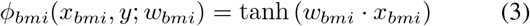

where *w*_*age*_ and *w*_*bmi*_ are the corresponding weights. We employed the age groups with an interval of 10 rather than the exact ages to alleviate the effect of age variability because age is an extremely sensitive risk factor for most diseases [28], [29] which may dominate the risk prediction.

For one-hot encoding, we assigned a learnable vector 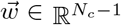 (*N*_*c*_ = the number of categories in the factor) to weight the variable. For example, we encoded the one-hot encoding of sex and cheese intake as below:

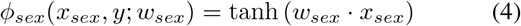

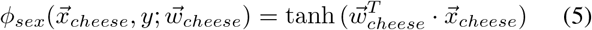

where *w*_*sex*_ *∈* ℝ^1^ and 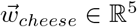 (*N*_*c*_ = 2 in sex; *N*_*c*_ = 6 in cheese intake) are the corresponding weights.

The log-linear model was then applied to aggregate and transform all encoding functions *ϕ* into a probabilistic distribution, which can be formulated as below:

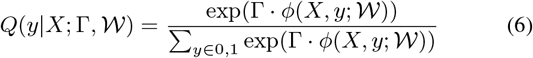

where Γ is the weight vector of encoding functions for each non-genetic factor. Based on the above examples of variable encodings (**Eq. 2, 3, 4, and 5**), the term Γ · *ϕ*(*X, y; 𝒲*) is equivalent to *γ*_*age*_ · *ϕ*_*age*_ + *γ*_*bmi*_ · *ϕ*_*bmi*_ + *γ*_*sex*_ · *ϕ*_*sex*_ + *γ*_*cheese*_ · *ϕ*_*cheese*_ (Γ = *γ*_*age*_, *γ*_*bmi*_, *γ*_*sex*_, *γ*_*cheese*_, 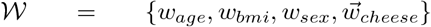, and 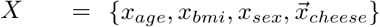). Similarly, we adopted cross-entropy as the loss function to optimize the log-linear model.

### E. Object function and risk score calculation

We applied posterior regularization [17], [30] technique to make *P* (**Eq. 1**) approximating *Q* (**Eq. 6**) by minimizing the KL-divergence between them. The KL loss term is formulated as below:

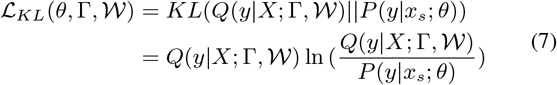

The objective function for global optimization consists of three loss terms, which can be calculated as below:

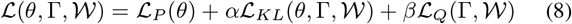

where *α* and *β* are two hyperparameters that can be tuned. *ℒ*_*P*_ *(θ)* and *ℒ*_*Q*_(Γ, 𝒲) are the cross-entropy loss of *P* (**Eq. 1**) and *Q* (**Eq. 6**), respectively. The final output of PRSIMD (*ŷ*) is formulated as below:

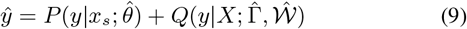

where 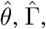 and 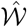 represent the estimated parameters learned from the training set.

### F. Model training and evaluation

PRSIMD was trained by Adam optimizer [31] with a mini-batch size of 128. We applied grid search ranging in {1, 0.1, 0.01, 0.001 to tune *α* and *β*. The model with the optimal loss on the validation set was selected to be evaluated on the test set (the right panel of **Fig. 1D**). We evaluated the predictive power of PRSIMD using AUROC, AUPRC (area under precision-recall curve), and RR (relative risk). For a given percentile X, RRs were calculated by comparing the ratios of incidence cases and total participants with prediction scores higher or lower than X. We utilized ”fmsb” [32] to calculate RRs and their confidence intervals.

## III. Results

### A. Effect sizes of the non-genetic factors

As shown in **Fig. 2A**, we found the forced vital capacity (OR=0.441, CI=[0.425, 0.458]) was the most significant factor associated with the risk of CAD negatively in physical measures. For the influence of lifestyles on CAD, increasing walking pace, cheese intake, and poultry intake was helpful to reduce the risk of CAD (**Fig. 2B**). The steady average (OR=0.153, CI=[0.135, 0.172]) and brisk (OR=0.083, CI=[0.073, 0.094]) walking paces had the lower risk of CAD compared with the slow walking paces. The light-to-vigorous cheese intake were significantly associated with the lower risk of CAD compared with never eating cheese. For example, eating greater or equal to once of cheese per day had a small OR value of 0.020 (CI=[0.015, 0.027]). The poultry intake frequency of 2-4 times per week was the most healthy frequency compared to the other frequencies of eating poultry, having a lower risk of CAD (OR=0.235, CI=[0.202, 0.272]) compared with never eating poultry. These three lifestyles were also proven to reduce the risk of CAD in previous studies (walking pace [33]; cheese intake [34]; and poultry intake [35]). Besides, the tobacco smoking on most or all day significantly increased the risk of CAD compared with no tobacco smoking in the present, and the other lifestyles had weak impacts on the disease risk (**Fig. 2B**).

**Fig. 2.**
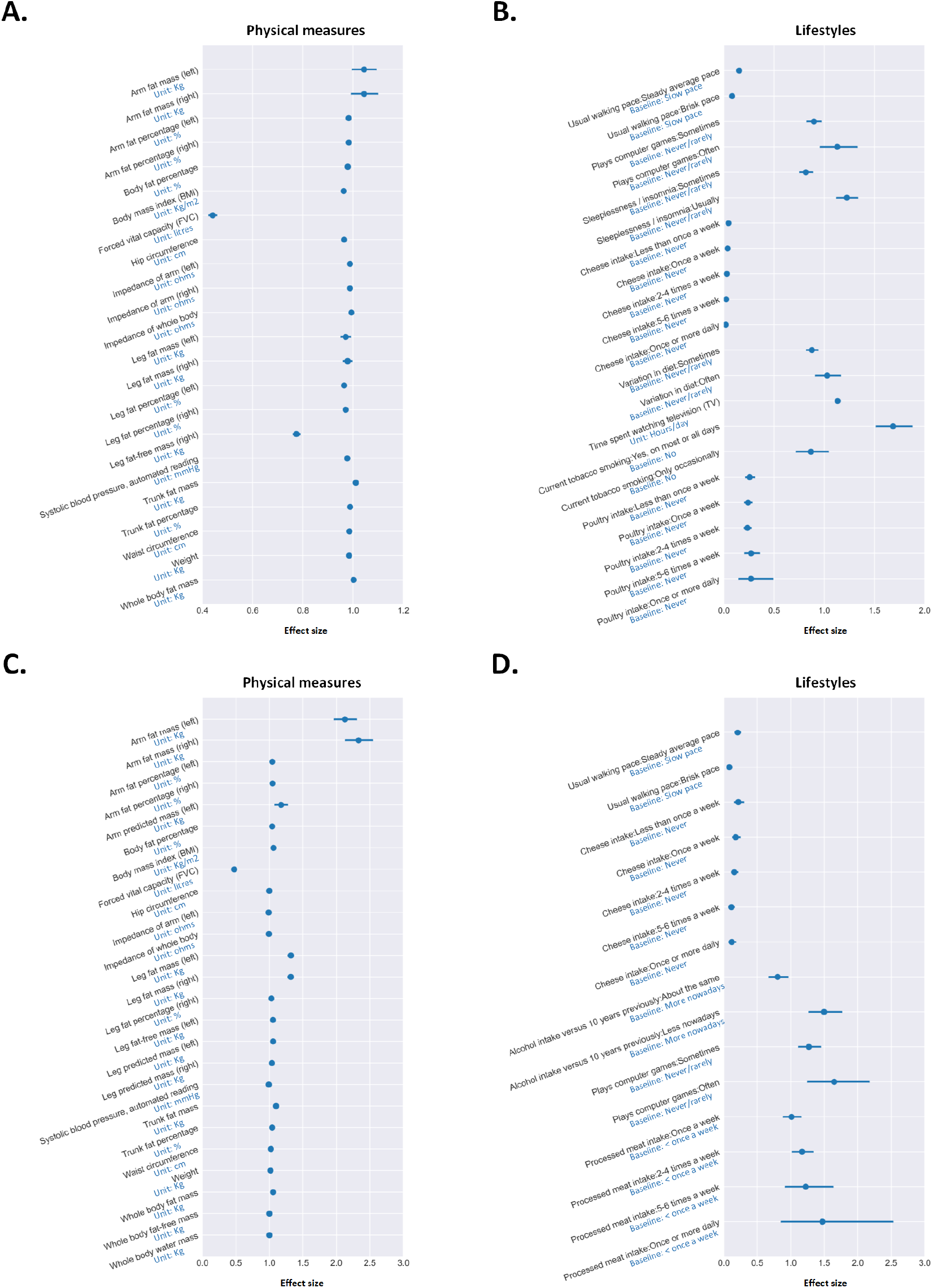
The effect sizes (dot) and 95% confidence intervals (line) of physical measures and lifestyles on CAD (**A** and **B**) and T2D (**C** and **D**). We calculated odds ratio as effect size for each non-genetic factor using the *L*1 penalized logistic regressions aggregated with age and sex. The odds ratio greater (less) than 1 indicates that the factor and the disease risk are positively (negatively) correlated.

As for the influence of physical measures on T2D, the higher value of arm (left: OR=2.127, CI=[1.968, 2.299]; right: OR=2.330, CI=[2.137, 2.540]) fat mass remarkably increased the disease risk. On the contrary, the higher value of forced vital capacity (OR=0.474, CI=[0.445, 0.504]) reduced the risk of T2D (**Fig. 2C**). As shown in **Fig. 2D**, increasing walking pace and cheese intake appropriately can reduce the risk of T2D, which was consistent with the trends of these two factors on the risk of CAD. Besides, the higher intake of processed meat relatively increased the risk of T2D. The increase in alcohol intake over the previous 10 years and the playing computer games frequently also contributed to the risk of T2D.

### B. Predictive performance on CAD and T2D

We evaluated PRSIMD and the other tools on CAD (8,747 cases) and T2D (2,838 cases) using over 1 million SNPs (**Table I**) and the selected non-genetic factor. We tried different tools to calculate PRSs, which were integrated into PRSIMD, and annotated them in the bracket (e.g., PRSIMD (LDpred2)). The predicted score percentiles were shown in the boxplots. As shown in **Fig. 3A-B**, PRSIMD remarkably improved disease risk prediction compared to the corresponding PRS method by introducing non-genetic factors. PRSIMD (LDpred2) achieved a best AUROC (AUPRC) of 0.760 (0.738) in predicting the risk of CAD (**Fig. 3A**), whereas the best predictive performance on T2D was PRSIMD (P+T) with an AUROC (AUPRC) of 0.846 (0.830). We observed that the performances of PRSIMD were consistent by incorporating different PRS methods. Besides, we extended CRS [15] to integrate the factors selected by MR causality inference (**Fig. 1C**) for disease risk prediction, and observed that PRSIMD also outperformed CRS (**Fig. 3C-D**). PRSIMD increased average AUROC of 0.021 and 0.056 on CAD and T2D compared to CRS, respectively.

**Fig. 3.**
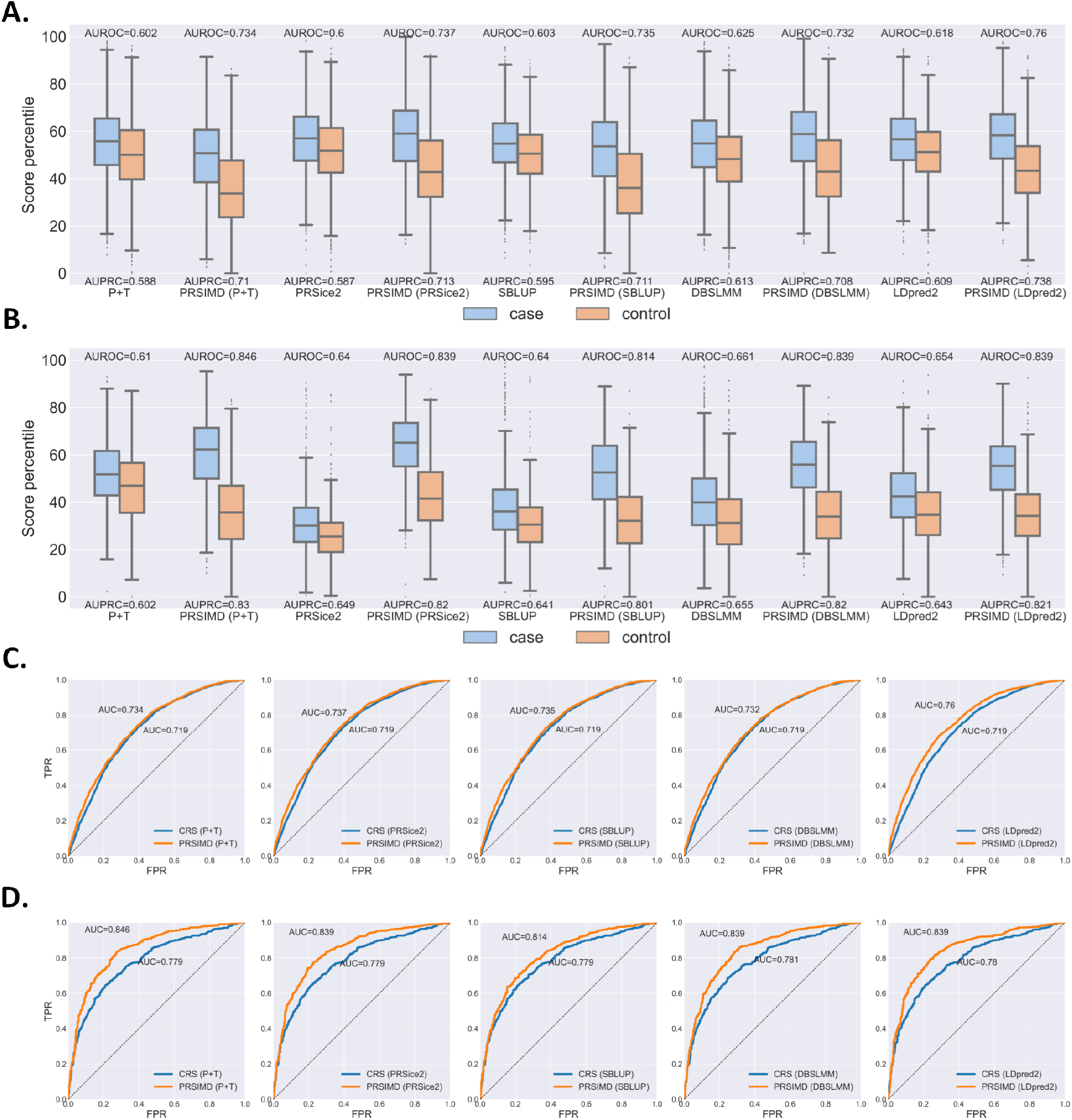
The predictive performance on CAD and T2D. **A-B**. The boxplots of the predicted score percentiles of CAD (**A**) and T2D (**B**) on five PRS methods and PRSIMD based on the corresponding one. The AUROCs and AUPRCs were shown over and under the boxplots of the corresponding methods, respectively. **C-D**. The ROCs of CRS and PRSIMD with five PRS methods.

### C. Relative risk on CAD and T2D

We defined the high-risk score percentile as top 20% (10%, 5%, 1%, and 0.5%) of the score distribution [20] to investigate the RRs based on the predicted scores of PRSIMD (LDpred2) on the test sets. The remaining scores under the cut-off of high-risk scores were used as the corresponding reference group. As shown in **Table. II**, the RRs (1.78 to 1.85 on CAD; 1.82 to 2.23 on T2D) were narrowly fluctuated among the different high-risk score definitions because we used the same amount of cases and controls on the test sets. PRSIMD (LDpred2) identified the top 20% had 1.85-fold and 2.23-fold risk on CAD and T2D compared to the reference groups, respectively.

**TABLE II.**
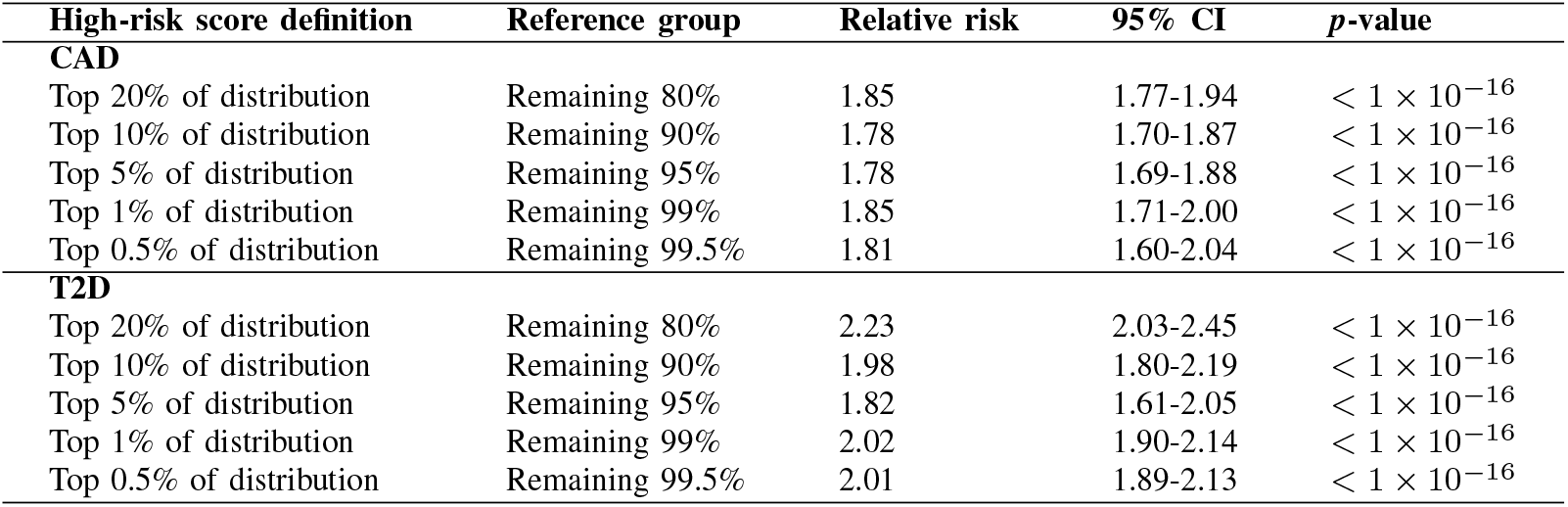
Prevalence of the high-risk scores

## IV. Discussion

PRS has become an effective tool to predict disease risk using an individual’s risk genetic variants. The non-genetic factors also significantly affect the risk of specific diseases, which were commonly neglected by the current tools. In this paper, we developed PRSIMD that attempted to integrate PRS and diverse non-genetic factors for disease risk prediction. We applied MR analysis to automatically choose disease causal risk factors from physical measures and lifestyles instead of selecting factors manually. These causal non-genetic factors can not only be used for disease risk prediction but also become potential interventions to prevent individuals from diseases. For example, high-risk individuals with CAD and T2D could consider increasing their cheese intake frequency and walking pace in daily life (**Fig. 2B, D**). Moreover, PRSIMD can flexibly incorporate non-genetic factors with various data types by constructing the encoding functions. The function tanh also can process the non-linear relationships between factors.

Besides, deep learning may be a feasible approach to incorporate different factors due to its powerful ability of data representation. DL-PRS [36] and DeepPRS [37] are two deep learning models proposed recently, utilizing fully connected neural networks and recurrent neural networks to calculate PRS. Their works proved that deep learning models could be applied to the genetic variants for risk prediction but required stringent SNP selection. We may attempt to apply deep learning techniques to integrate both genetic and non-genetic factors in the future.

## Author’s contributions

LZ conceived the study; YX implemented PRSIMD and analyzed the results. YX, CHW, ZML conducted the experiments. YX wrote the article. LZ, YPC, OZY, and APL reviewed the paper. All authors read and approved the final manuscript.

## Acknowledgements

This research is partially supported by TCFS (GHX/133/20SZ), Hong Kong Research Grant Council Early Career Scheme (HKBU 22201419), Guangdong Basic and Applied Basic Research Foundation (No. 2021A1515012226) and SZVUP Special Fund Project (2021Szvup135). This research conducted using the UK Biobank Resource under Application 60434.

## Code available

The codes of PRSIMD are available at: https://github.com/yuxu-1/PRSIMD

## References

[1] S. W. Choi, T. S.-H. Mak, and P. F. O’Reilly, “Tutorial: a guide to performing polygenic risk score analyses,” Nature protocols, vol. 15, no. 9, pp. 2759–2772, 2020.

[2] C. Wang, J. Zhang, X. Zhou, and L. Zhang, “A comprehensive investigation of statistical and machine learning approaches for predicting complex human diseases on genomic variants,” bioRxiv, 2022.

[3] B. J. Vilhjálmsson, J. Yang, H. K. Finucane, A. Gusev, S. Lindström, S. Ripke, G. Genovese, P.-R. Loh, G. Bhatia, R. Do et al., “Modeling linkage disequilibrium increases accuracy of polygenic risk scores,” The american journal of human genetics, vol. 97, no. 4, pp. 576–592, 2015.

[4] S. W. Choi and P. F. O’Reilly, “PRSice-2: Polygenic Risk Score software for biobank-scale data,” GigaScience, vol. 8, no. 7, 07 2019.

[5] M. R. Robinson, A. Kleinman, M. Graff, A. A. Vinkhuyzen, D. Couper, M. B. Miller, W. J. Peyrot, A. Abdellaoui, B. P. Zietsch, I. M. Nolte et al., “Genetic evidence of assortative mating in humans,” Nature Human Behaviour, vol. 1, no. 1, pp. 1–13, 2017.

[6] S. Yang and X. Zhou, “Accurate and scalable construction of polygenic scores in large biobank data sets,” The American Journal of Human Genetics, vol. 106, no. 5, pp. 679–693, 2020.

[7] F. Privé, J. Arbel, and B. J. Vilhjálmsson, “Ldpred2: better, faster, stronger,” Bioinformatics, vol. 36, no. 22-23, pp. 5424–5431, 2020.

[8] T. Ge, C.-Y. Chen, Y. Ni, Y.-C. A. Feng, and J. W. Smoller, “Polygenic prediction via bayesian regression and continuous shrinkage priors,” Nature communications, vol. 10, no. 1, pp. 1–10, 2019.

[9] M. Nikpay, A. Goel, H.-H. Won, L. M. Hall, C. Willenborg, S. Kanoni, D. Saleheen, T. Kyriakou, C. P. Nelson, J. C. Hopewell et al., “A comprehensive 1000 genomes-based genome-wide association meta-analysis of coronary artery disease,” Nature genetics, vol. 47, no. 10, p. 1121, 2015.

[10] R. A. Scott, L. J. Scott, R. Mägi, L. Marullo, K. J. Gaulton, M. Kaakinen, N. Pervjakova, T. H. Pers, A. D. Johnson, J. D. Eicher et al., “An expanded genome-wide association study of type 2 diabetes in europeans,” Diabetes, vol. 66, no. 11, pp. 2888–2902, 2017.

[11] M. Inouye, G. Abraham, C. P. Nelson, A. M. Wood, M. J. Sweeting, F. Dudbridge, F. Y. Lai, S. Kaptoge, M. Brozynska, T. Wang et al., “Genomic risk prediction of coronary artery disease in 480,000 adults: implications for primary prevention,” Journal of the American College of Cardiology, vol. 72, no. 16, pp. 1883–1893, 2018.

[12] N. Mavaddat, K. Michailidou, J. Dennis, M. Lush, L. Fachal, A. Lee, J. P. Tyrer, T.-H. Chen, Q. Wang, M. K. Bolla et al., “Polygenic risk scores for prediction of breast cancer and breast cancer subtypes,” The American Journal of Human Genetics, vol. 104, no. 1, pp. 21–34, 2019.

[13] J. W. O’Sullivan, A. Shcherbina, J. M. Justesen, M. Turakhia, M. Perez, H. Wand, C. Tcheandjieu, S. L. Clarke, M. A. Rivas, and E. A. Ashley, “Combining clinical and polygenic risk improves stroke prediction among individuals with atrial fibrillation,” Circulation: Genomic and Precision Medicine, vol. 14, no. 3, p. e003168, 2021.

[14] Y. Shieh, D. Hu, L. Ma, S. Huntsman, C. C. Gard, J. W. Leung, J. A. Tice, C. M. Vachon, S. R. Cummings, K. Kerlikowske et al., “Breast cancer risk prediction using a clinical risk model and polygenic risk score,” Breast cancer research and treatment, vol. 159, no. 3, pp. 513–525, 2016.

[15] A. Moldovan, Y. Y. Waldman, N. Brandes, and M. Linial, “Body mass index and birth weight improve polygenic risk score for type 2 diabetes,” Journal of personalized medicine, vol. 11, no. 6, p. 582, 2021.

[16] K. Ganchev, J. Graça, J. Gillenwater, and B. Taskar, “Posterior regularization for structured latent variable models,” The Journal of Machine Learning Research, vol. 11, pp. 2001–2049, 2010.

[17] F. Ma, J. Gao, Q. Suo, Q. You, J. Zhou, and A. Zhang, “Risk prediction on electronic health records with prior medical knowledge,” in Proceedings of the 24th ACM SIGKDD International Conference on Knowledge Discovery & Data Mining, 2018, pp. 1910–1919.

[18] C. Sudlow, J. Gallacher, N. Allen, V. Beral, P. Burton, J. Danesh, P. Downey, P. Elliott, J. Green, M. Landray et al., “Uk biobank: an open access resource for identifying the causes of a wide range of complex diseases of middle and old age,” PLoS medicine, vol. 12, no. 3, p. e1001779, 2015.

[19] O. O. Yavorska and S. Burgess, “Mendelianrandomization: an r package for performing mendelian randomization analyses using summarized data,” International journal of epidemiology, vol. 46, no. 6, pp. 1734–1739, 2017.

[20] A. V. Khera, M. Chaffin, K. G. Aragam, M. E. Haas, C. Roselli, S. H. Choi, P. Natarajan, E. S. Lander, S. A. Lubitz, P. T. Ellinor et al., “Genome-wide polygenic scores for common diseases identify individuals with risk equivalent to monogenic mutations,” Nature genetics, vol. 50, no. 9, pp. 1219–1224, 2018.

[21] B. Elsworth, M. Lyon, T. Alexander, Y. Liu, P. Matthews, J. Hallett, P. Bates, T. Palmer, V. Haberland, G. D. Smith et al., “The mrc ieu opengwas data infrastructure,” BioRxiv, 2020.

[22] G. Hemani, J. Zheng, B. Elsworth, K. H. Wade, V. Haberland, D. Baird, C. Laurin, S. Burgess, J. Bowden, R. Langdon et al., “The mrbase platform supports systematic causal inference across the human phenome,” elife, vol. 7, 2018.

[23] F. Yang, T. Hu, S. Chen, K. Wang, Z. Qu, and H. Cui, “Low intelligence predicts higher risks of coronary artery disease and myocardial infarction: Evidence from mendelian randomization study,” Frontiers in genetics, vol. 13, 2022.

[24] J. R. Staley, J. Blackshaw, M. A. Kamat, S. Ellis, P. Surendran, B. B. Sun, D. S. Paul, D. Freitag, S. Burgess, J. Danesh et al., “Phenoscanner: a database of human genotype–phenotype associations,” Bioinformatics, vol. 32, no. 20, pp. 3207–3209, 2016.

[25] G. Hemani, J. Zheng, B. Elsworth, K. Wade, D. Baird, V. Haberland, C. Laurin, S. Burgess, J. Bowden, R. Langdon, V. Tan, J. Yarmolinsky, H. Shibab, N. Timpson, D. Evans, C. Relton, R. Martin, G. Davey Smith, T. Gaunt, P. Haycock, and The MR-Base Collaboration, “The mr-base platform supports systematic causal inference across the human phenome,” eLife, vol. 7, p. e34408, 2018. [Online]. Available: https://elifesciences.org/articles/34408

[26] P. Van Der Harst and N. Verweij, “Identification of 64 novel genetic loci provides an expanded view on the genetic architecture of coronary artery disease,” Circulation research, vol. 122, no. 3, pp. 433–443, 2018.

[27] P.-T. De Boer, D. P. Kroese, S. Mannor, and R. Y. Rubinstein, “A tutorial on the cross-entropy method,” Annals of operations research, vol. 134, no. 1, pp. 19–67, 2005.

[28] M. Kamińska, T. Ciszewski, K. Łopacka-Szatan, P. Miotła, and E. Starosławska, “Breast cancer risk factors,” Przeglad menopauzalny= Menopause review, vol. 14, no. 3, p. 196, 2015.

[29] A. Sonnenberg, “Age distribution of ibd hospitalization,” Inflammatory bowel diseases, vol. 16, no. 3, pp. 452–457, 2010.

[30] J. Zhang, Y. Liu, H. Luan, J. Xu, and M. Sun, “Prior knowledge integration for neural machine translation using posterior regularization,” arXiv preprint 1811.01100, 2018.

[31] D. P. Kingma and J. Ba, “Adam: A method for stochastic optimization,” arXiv preprint 1412.6980, 2014.

[32] M. Nakazawa and M. M. Nakazawa, “Package ‘fmsb’,” See https://cran.r-project.org/web/packages/fmsb/fmsb.pdf, 2019.

[33] E. Stamatakis, P. Kelly, T. Strain, E. M. Murtagh, D. Ding, and M. H. Murphy, “Self-rated walking pace and all-cause, cardiovascular disease and cancer mortality: individual participant pooled analysis of 50 225 walkers from 11 population british cohorts,” British Journal of Sports Medicine, vol. 52, no. 12, pp. 761–768, 2018.

[34] M.-J. Hu, J.-S. Tan, X.-J. Gao, J.-G. Yang, and Y.-J. Yang, “Effect of cheese intake on cardiovascular diseases and cardiovascular biomarkers,” Nutrients, vol. 14, no. 14, p. 2936, 2022.

[35] K. Hoffmann, B.-C. Zyriax, H. Boeing, and E. Windler, “A dietary pattern derived to explain biomarker variation is strongly associated with the risk of coronary artery disease,” The American journal of clinical nutrition, vol. 80, no. 3, pp. 633–640, 2004.

[36] S. Huang, X. Ji, M. Cho, J. Joo, and J. Moore, “Dl-prs: a novel deep learning approach to polygenic risk scores,” 2021.

[37] J. Peng, J. Li, R. Han, Y. Wang, L. Han, J. Peng, T. Wang, J. Hao, X. Shang, and Z. Wei, “A deep learning-based genome-wide polygenic risk score for common diseases identifies individuals with risk,” medRxiv, 2021.

